# Structural remodeling of *Coxiella burnetii* during its biphasic developmental cycle revealed by cryo-electron tomography

**DOI:** 10.1101/2022.08.23.505044

**Authors:** Mohammed Kaplan, Doulin C. Shepherd, Naveen Vankadari, Ki Woo Kim, Charles L. Larson, Przemysław Dutka, Paul A. Beare, Edward Krzymowski, Robert A. Heinzen, Grant J. Jensen, Debnath Ghosal

**Affiliations:** Division of Biology and Biological Engineering, California Institute of Technology, Pasadena, CA 91125, USA; Department of Biochemistry and Pharmacology, Bio21 Molecular Science and Biotechnology Institute, The University of Melbourne, Melbourne, VIC, Australia; School of Ecology and Environmental System, Kyungpook National University, Sangju, Korea; Coxiella Pathogenesis Section, Laboratory of Bacteriology, Rocky Mountain Laboratories, National Institute of Allergy and Infectious Diseases, National Institutes of Health, Hamilton, Montana, United States of America; Division od Chemistry and Chemical Engineering, California Institute of Technology, 1200 California Boulevard, Pasadena, CA 91125, USA; Department of Physics and Astronomy, Brigham Young University, Provo, UT 84604, USA; Department of Chemistry and Biochemistry, Brigham Young University, Provo, UT 84604, USA; ARC Centre for Cryo-electron Microscopy of Membrane Proteins, Bio21 Molecular Science and Biotechnology Institute, University of Melbourne, Parkville, Victoria, Australia

**Keywords:** *Coxiella burnetii*, intracellular pathogen, biphasic life cycle, LCV, TCV, SCV, Dot/Icm T4SS, cryo-electron tomography

## Abstract

*Coxiella burnetii* is an obligate zoonotic bacterium that targets macrophages to cause a disease known as Q fever. It has a biphasic developmental lifecycle where the extracellular and metabolically inactive small cell variant (SCV) transforms, under host acidic environment, into the vegetative large cell variant (LCV). However, the details about the morphological and structural changes that accompany this biphasic cycle are still lacking. Here, we used cryo-electron tomography to image the different cell variants of *C. burnetii* grown either under axenic conditions in different pH or purified directly from host cells revealing the major developmental, morphological and structural transitions. We show that SCVs are characterized by equidistant stacks of inner membrane that presumably allow a smooth transition to LCV, a transition coupled with the expression of the Dot/Icm type IVB secretion system (T4BSS). A class of T4BSS particles were associated with extracellular densities including a tubular structure possibly involved in host interaction or effector delivery. Also, SCVs and cells in the transition state contained spherical multilayered membrane structures of different sizes and locations suggesting that they are not related to a sporulation process as once assumed.

## INTRODUCTION

*Coxiella burnetii* is a highly infectious wide-ranging zoonotic pathogenic bacterium and the causative agent of human Q fever, a severe and debilitating form of influenza-like illness (1–3). *C. burnetii* exhibits a unique biphasic developmental cycle of metabolically active and replicating (exponential phase) large cell variant (LCV) present inside the host, to nonreplicating (stationary phase) dormant small cell variant (SCV) that can survive outside the host cell (4–7). Each variant has distinct morphological features; while SCVs are characterized by their rod-shape, shorter length (0.2 to 0.6 μm), dense periplasmic space and condensed DNA, LCVs are pleomorphic in shape, with length exceeding 1 μm, and exhibit relatively sparse periplasmic space and dispersed DNA (4–6,8). The SCVs are highly stable in external environments (outside the host cell) and recalcitrant to physical and chemical stresses (4,9–11). The primary mode of human infection is inhalation of mainly SCVs through contaminated aerosol (12). *C. burnetii* SCVs predominantly target primary mononuclear phagocytes (e.g., alveolar macrophages) to begin the intracellular biphasic life cycle (13,14). Following internalization, *C. burnetii* establishes a specialized membrane-bound replicative niche known as the *Coxiella* containing vacuole (CCV) which matures into an autophagolysosome-like compartment with an acidified lumen (pH~4.75), acid hydrolases, cationic peptides, and lysosomal markers such as LAMP-1 and CD63 (LAMP-3) (15,16). The acidification of the CCV upregulates metabolic activity and gene expression in SCVs leading to a transition to the metabolically active LCVs (17–19).

The maintenance and maturation of the CCV is largely mediated by the activity of the *C. burnetii* Dot/Icm (defective in organelle trafficking/intracellular multiplication) type IV secretion system (T4SS) that delivers more than 130 unique effector proteins into the host cytosol (20–25). The bacterial T4SSs are multimegadalton molecular machines that span through the bacterial cell envelope and are involved in transport of nucleoprotein complexes as well as proteins across the cellular envelope (26). Based on genetic composition and phylogenetic analysis, they are classified into two major classes, T4ASS and T4BSS (27,28). The prototype for the T4A and T4BSS are the *Agrobacterium tumefaciens* and *Legionella pneumophila* T4SSs, respectively. The *C. burnetii* T4SS, which belongs to the T4BSS, is phylogenetically very closely related to the *L. pneumophila* T4BSS and comprises at least 23 homologues of the ~30 *L. pneumophila dot/icm* genes.

*C. burnetii* T4BSS effector proteins subvert multiple host cellular pathways such as autophagic, secretory, and endolysosomal trafficking and aid biogenesis of the CCV to facilitate intracellular replication of *C. burnetii* (29–32). Earlier studies suggested that the transition from SCV to LCV inside the CCV is correlated with the upregulation of T4BSS expression and activity (8,18,33,34). Intriguingly, under *in vitro* and axenic (host cell-free) growth conditions, *C. burnetii* slowly differentiates from LCVs to SCVs; this transition is evident after 10 days, and the majority of LCVs convert to SCVs after 21 days (35).

Despite decades of research, the structural changes and remodeling that occur in *C. burnetii* during its biphasic cycle from LCV to SCV and *vice versa* remain elusive in part due to the lack of high-resolution *in situ* imaging of this obligate pathogen. While recent advances in single particle cryo-electron microscopy (cryo-EM) and cryo-electron tomography (cryo-ET) methods have enabled the investigation of T4SSs in great detail in various bacterial species (36–41), the molecular architecture, localization and developmental regulation of the *C. burnetii* T4BSS with respect to its biphasic cycle remain poorly understood. Moreover, previous studies using conventional transmission electron microscopy (TEM) suggested the formation of “spore-like” structures, together with membrane remodeling, at the transition from LCV to SCV. However, the existence of such structures and remodeling remain controversial as sample preparation (fixation and dehydration) and staining in conventional TEM disrupts membranes. Here, we used cryo-ET to visualize *C. burnetii* LCVs and SCVs in axenic (host cell-free) growth conditions as well as host-derived *C. burnetii* variants at molecular resolution. Our comprehensive cryo-ET analyses captured structural changes in cellular morphology and intricate membrane dynamics associated with LCV to SCV transition and vice versa. In addition, *in situ* structural analyses of the *C. burnetii* Dot/Icm T4BSS under different developmental conditions provided insights into tight regulation of the Dot/Icm T4BSS expression and activity.

## RESULTS

### Molecular architecture of the intact *C. burnetii* T4BSS and its regulation during the biphasic developmental cycle

To reveal the molecular architecture of the *C. burnetii* T4BSS and its regulation during different developmental stages, we used cryo-ET to image frozen hydrated *C. burnetii* cells (both SCVs and LCVs) grown in host cell-free second-generation acidified citrate cysteine media (ACCM-2). Earlier studies showed that the biphasic developmental transition and the growth kinetics and viability of *C. burnetii* can be recapitulated in ACCM-2 media (35). For example, when grown in axenic ACCM-2 media, nearly all SCVs transition to LCVs after 4 days, and then SCVs start appearing around day 10 and become the predominant cell variant again around day 14 (35). After 21 days in ACCM-2, nearly all LCVs transition to SCVs (35). These transition and growth kinetic of *C. burnetii* are very similar to those occurring inside a host cell such as Vero (African green monkey kidney fibroblasts) cells (18,35). Hence, we decided to image *C. burnetii* grown in ACCM-2 medium for 5 and 14 days to capture the morphological and structural signatures of SCV to LCV transition and vice versa. This approach is advantageous for cryo-ET imaging where the small size of *C. burnetii* makes them suitable for direct imaging, unlike thicker cells (e.g., eukaryotic cells) which must be thinned in order to become amenable for tomography data collection.

Cryo-ET imaging of the day 5 axenic culture showed that ~62% of the cells are LCVs (cell size > 800 nm), ~35% cells are in their transition state (transition state cell variant (TCV), 600-800 nm) and only 3% cells were SCVs (< 600 nm) (SI Fig. 1). Compared to SCVs, LCVs exhibited relatively sparse periplasmic space, dispersed DNA, and contained many “Wi-Fi” shaped structures spanning across the cell envelope (Fig. 1 A-B). These structures, ~4 per cell, were composed of two major densities - an outer membrane (OM) associated layer and a lower periplasmic layer (Fig. 1C). In partially lysed cells, we occasionally observed clear top views of these particles that appeared to have two concentric rings with the outer ring diameter being ~40 nm (Fig. 1D). Due to their similarity to the T4BSS particles in *L. pneumophila* (37), we hypothesized that these particles are the T4BSS of *C. burnetii*. Accordingly, these particles were absent in 17 cryotomograms of a *C. burnetii* strain with a complete *dot/icm* knockout, confirming that they are T4BSS particles. In contrast, cryo-ET imaging of the day 15 axenic culture contained a mixed population of SCVs (19%), LCVs (37 %) and TCVs (44%), and here also T4BSS particles were present only in LCVs and TCVs but not in SCVs (SI Fig. 2). Our observation that the assembly and maintenance of the T4BSS apparatus is coupled to the developmental transition from SCV to LCV is consistent with previous studies (8,18,33,34).

**Figure 1:**
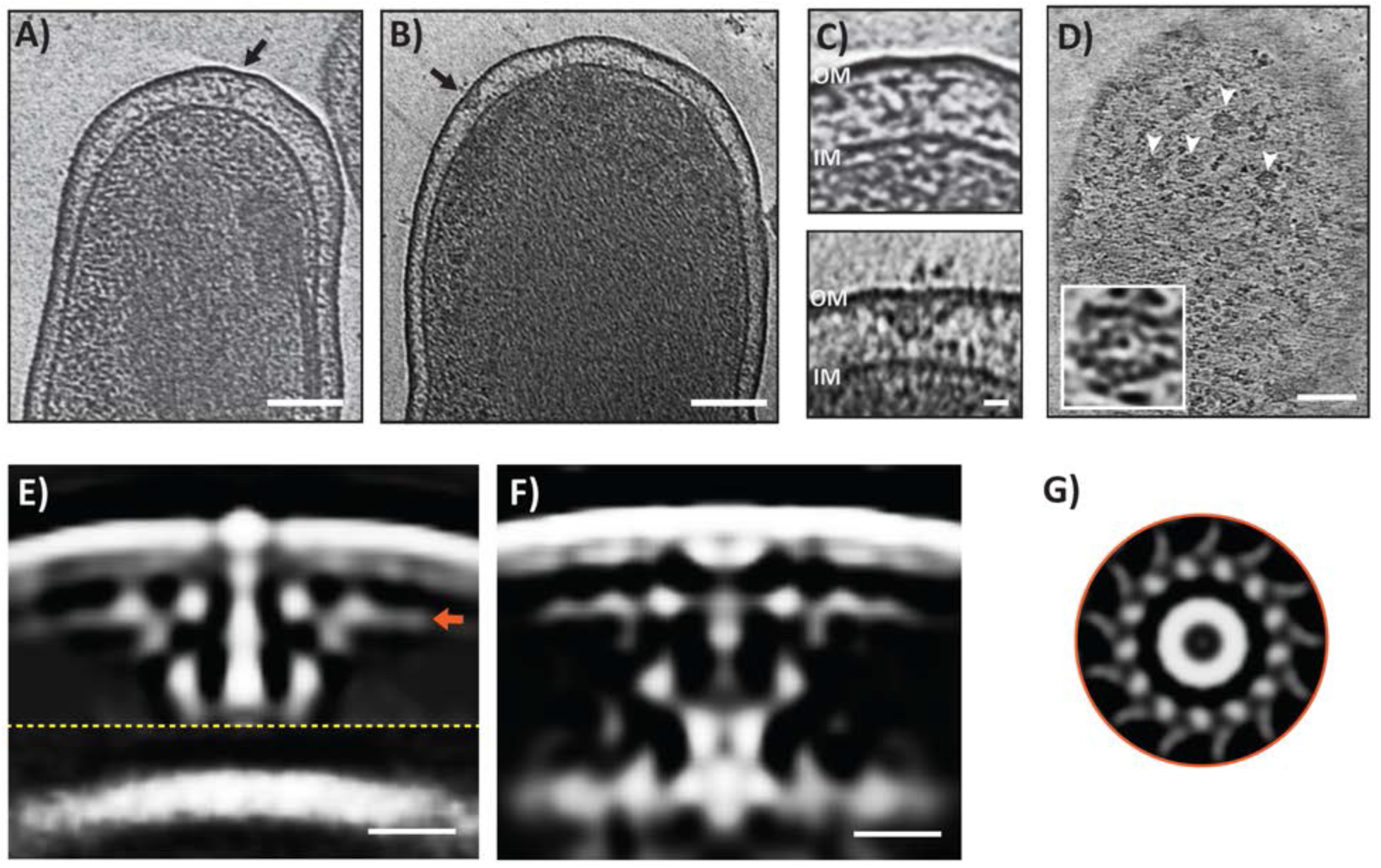
A&B) Slices through electron cryotomograms of *C. burnetii* cells highlighting the presence of T4SS particles (black arrows). Scale bar 100 nm. C) Enlargements of the T4SS particles shown in A (top panel) and B (lower panel). Scale bar 10 nm. D) A slice thorough an electron cryotomogram of a lysed *C. burnetii* cell illustrating the presence of multiple top views of T4SS particles (white arrows). Scale bar 100 nm. An enlargement of a top view of a T4SS particle is shown in the white-boxed area. E&F) Slices through the subtomogram averages of T4SS of *C. burnetii* at pH7.2 (left) and pH4.75 (right). Dashed yellow line indicates a composite of two averages obtained with different masks (see materials and methods) concatenated together. Scale bar 10 nm. G) A cross section through the T4SS OM complex (at the level indicated with the orange arrow in E) showing 13-fold symmetry. OM= outer membrane, IM= inner membrane.

**Figure 2:**
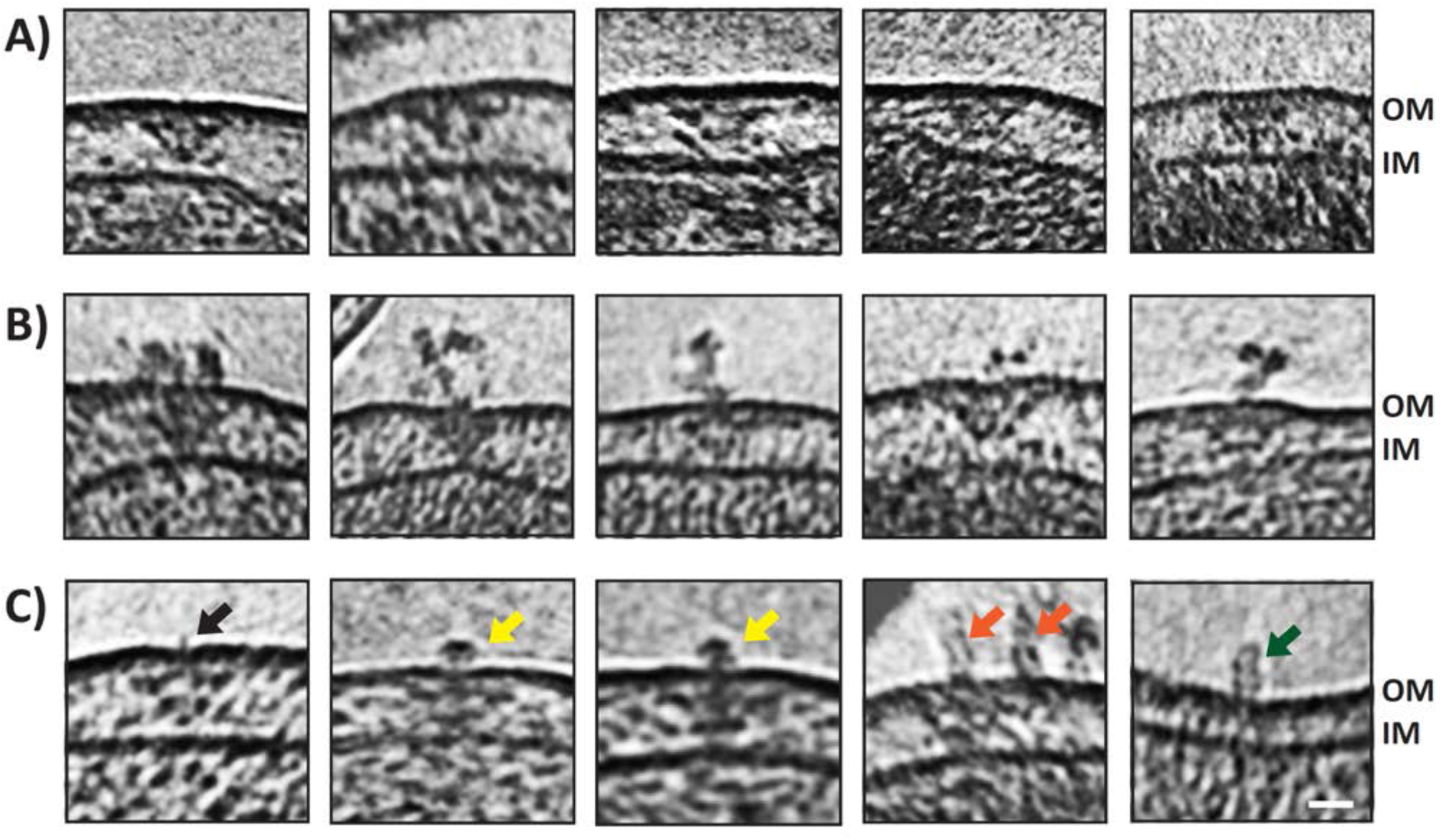
A) Slices through electron cryotomograms of *C. burnetii* cells indicating the presence of T4SS particles without any extracellular densities associated with them. B) Slices through electron cryotomograms of *C. burnetii* cells showing the presence of unstructured extracellular densities above the T4SS particles. C) Slices through electron cryotomograms of *C. burnetii* cells illustrating different types of extracellular densities associated with T4SS such as: short filament-like density (black arrow), crown-like density (yellow arrows), tubular densities (orange arrows). Green arrow point to tubular densities where no T4SS could be identified at their bases. Scale bar is 20 nm. OM= outer membrane, IM= inner membrane.

While the majority of T4BSS particles of *C. burnetii* were located at the cell poles, there were also a few (7 in 138 tomograms) that were positioned away from the cell poles (SI Fig. 3A). In our previous work on *L. pneumophila*, we observed only a few T4BSS particles located away from the cell pole in > 3,500 cryotomograms (42,43). However, similar to our previous observation in *L. pneumophila*, we identified T4BSS particles at the division plane in dividing *C. burnetii*, suggesting that they start assembling at the future new poles during the septation process (SI Fig. 3B).

**Figure 3:**
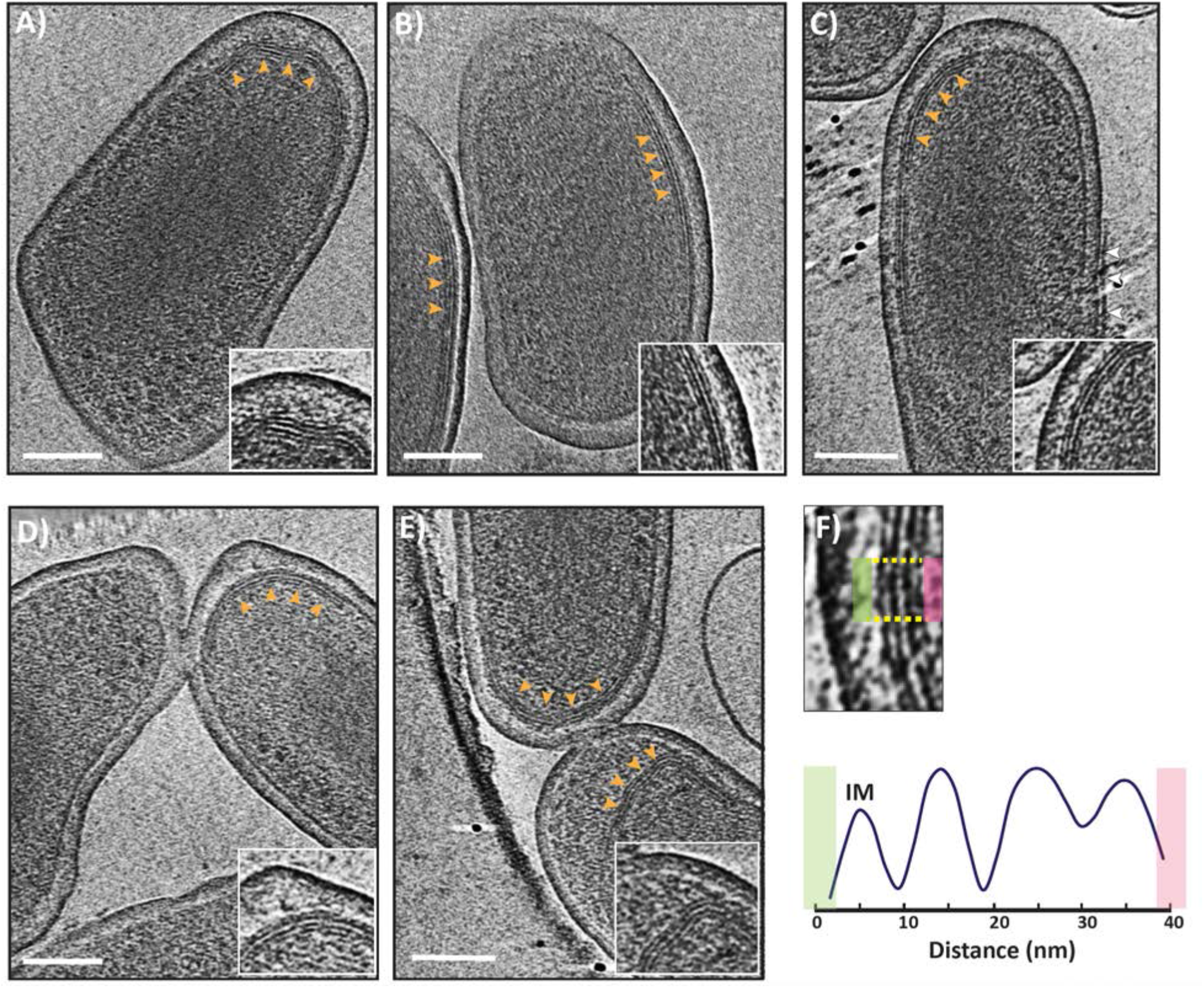
A-E) Slices through electron cryotomograms of *C. burnetii* cells (purified from infected vero cells 28 dpi) highlighting the presence of multilayered cytoplasmic membrane invaginations (orange arrows) with zoom-ins of these invaginations shown in the enlargements at the lower right corner of each panel. The white arrows in panel C point to a membrane of presumably the host vacuole still attached to the OM of *C. burnetii* cell. Scale bar is 100 nm. F) Lower panel: average density profile taken along the area indicated between the yellow lines in the upper panel highlighting the equidistant spacing of ~10 nm between the different layers of the inner membrane invaginations.

To decipher the molecular architecture of the *C. burnetii* T4BSS, we averaged 512 particles and generated a subtomogram average at a resolution of 2.5-4.5 nm (Fig. 1E and SI Fig. 4A). This average was generated from cells grown in ACCM-2 for 5 days and resuspended in phosphate buffer saline pH 7.2. Our initial average nicely resolved densities associated with the OM as well as the periplasmic complex; however, the inner membrane (IM) associated densities were missing. A closer look at the individual particles revealed that the distance between the OM and IM varied significantly in individual particles with most of them lacking clear densities associated with the IM (Fig. 1C and later in Fig. 2). Consequently, we generated two separate averages with different masks, one with a mask on the OM and one on the IM and juxtaposed them together to produce a composite average with a local resolution between 2.5-4.5 nm (Fig. 1E and SI Fig. 4A). This composite average showed marked structural similarity to the T4BSS of *L. pneumophila*, where both consisted of an OM complex, a periplasmic complex and a stem-like structure connecting the two (SI Fig. 5A-B). The OM complex showed a 13-fold symmetry and a maximum diameter of ~40 nm (Fig. 1G), and formed together with the periplasmic complex a secretion chamber similar to the *L. pneumophila* T4BSS system. This structural similarity between the two systems is not surprising given that there is a clear homology between at least 23 genes of the 30 T4BSS genes in *L. pneumophila* (44).

**Figure 4:**
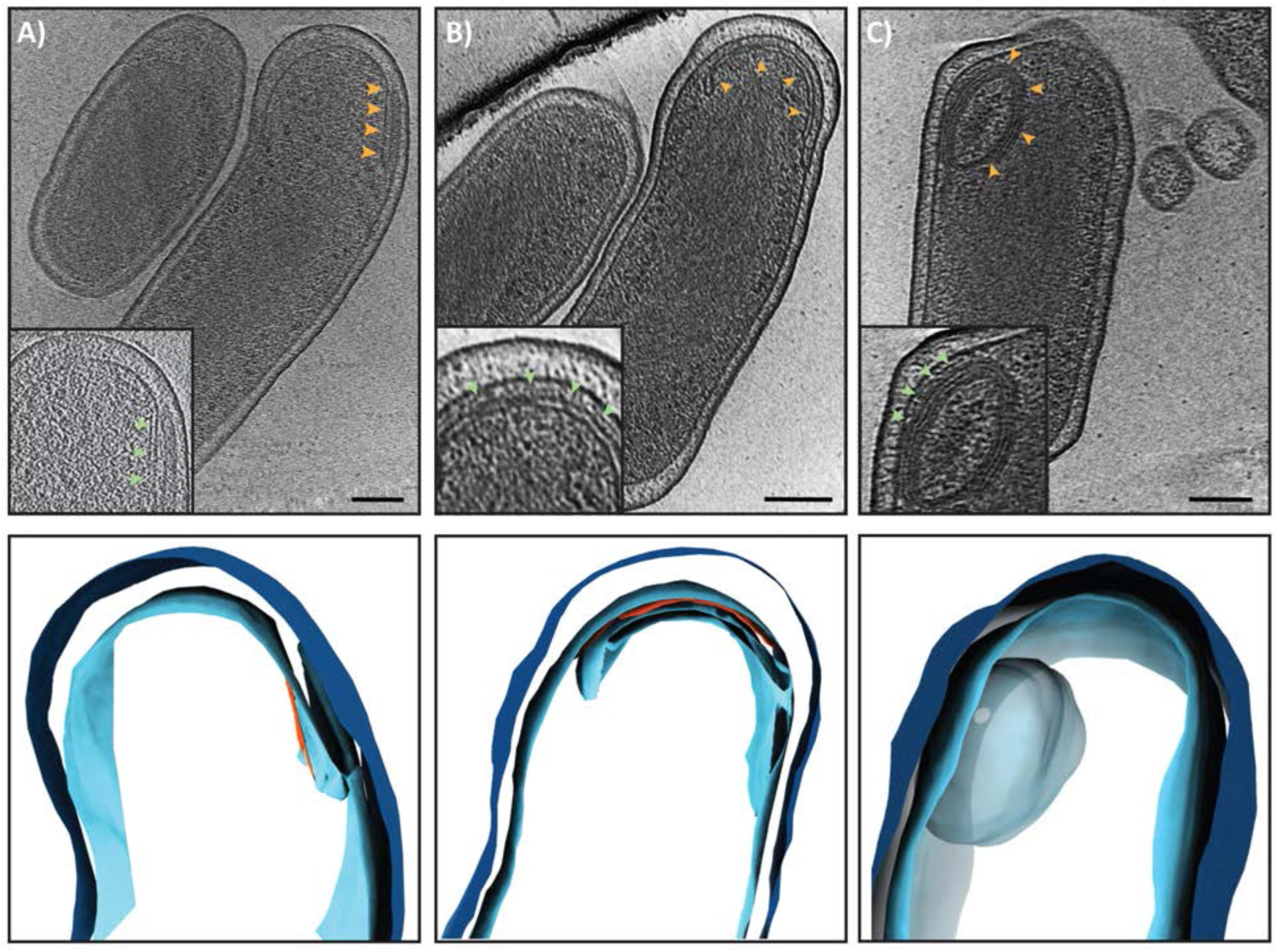
A-C) Top panels: Slices through electron cryotomograms of *C. burnetii* cells (grown in ACCM-2 for 15 days) illustrating different stages of inner membrane invaginations (orange arrows) with 3D segmentations of these stages shown in the lower panels. An early stage of IM invagination is shown in panel A where a protein-like density can be seen at one side of this invagination (red density in the segmentation). Panel B shows a long and curved IM invagination where a protein-like density can be seen between this invagination and the IM (red density in the segmentation, green arrow in inset). A whole multilayered vesicular structure can be seen in panel C. Note that it was difficult to trace the different layers in the tomogram hence it is shown as one large blob in the segmentation in the lower panel (See SI Movie 2). Scale bar is 100 nm.

Subsequently, we docked the recently-reported near-atomic resolution structure of the *L. pneumophila* T4BSS core complex (45) into our subtomogram average which allowed us to tentatively assign components that contribute to different densities in our structure. This fitting suggested that the OM core complex comprised DotC, DotD, DotF, DotH, DotG and DotK proteins while the periplasmic ring complex contains parts of DotG and DotH (SI Fig. 6). On the other hand, and despite several data processing trials, we could not resolve the cytoplasmic ATPase complex. A contemporaneous study on *C. burnetii* T4BSS used more than 7000 particles to generate an *in situ* average (46), but their structure also lacked the cytoplasmic complex suggesting that the *C. burnetii* T4BSS ATPase complex is either flexible or rarely associated. Earlier studies have shown that acidification of the CCV inside the host triggers the differentiation of SCVs into LCVs. This developmental transition has been linked to enhanced metabolic activities, higher gene expression and biogenesis and secretion of the *C. burnetii* T4BSS (8,18,33,34). Based on this, we hypothesized that under autolysosomal pH conditions (pH ~4.75), we might capture actively secreting T4BSSs in our cryotomograms. Therefore, we cultured *C. burnetii* cells in ACCM-2 media for 5 days and then resuspended them in a low pH media (citrate buffer saline pH ~4.75). Unlike the particles observed in the previous tomograms of cells at pH 7.2, individual particles in these tomograms of cells at pH 4.75 showed less variability in the distance between the OM and IM. We used 411 particles to produce a subtomogram average (with a single mask) at 2.5–5 nm resolution (Fig. 1F, SI Fig. 4B) which revealed a few structural differences (as suggested at the resolution of our structures) compared to the average at pH 7 including the existence of a secretion channel and density for the wing structures (as seen before in the *L. pneumophila* T4BSS, SI Fig. 5). These observations are consistent with the fact that the T4BSS is active in the acidic autolysosome environment and therefore acidic pH might have ‘primed’ these complexes.

Interestingly, ~17% of individual T4BSS particles showed extracellular densities associated with them at day 5 (both pH 4.75 and 7.2) and day 15 cultures. We classified these extracellular densities into 4 major categories: i) those with an amorphous shape, ii) thin filament-like iii) crown-like structures and iv) those with a rod or tubular-like form (Fig. 2A-C). However, some of these extracellular densities were also occasionally localized on the cell surfaces without an obvious T4BSS particle underneath them (Fig. 2C, right most). This could be either because they are relics after the T4BSSs are disassembled or they were secreted and then drifted away from T4BSS. Alternatively, it is also possible that they were secreted in a T4BSS-independent manner. The low number of particles with extracellular densities precluded the possibility of producing a decent subtomogram average to see if there are any structural changes in T4BSS associated with the presence of these extracellular densities.

Of the different types, the ‘tube-like’ densities were the most interesting as they are reminiscent of a T4ASS pilus filament. These tube-like filaments were ~8 nm wide, ~25 nm in length and showed flexibility. In our more than 3500 tomograms of *L. pneumophila* cells, we never saw T4BSS particles with any extracellular density (42,43). Furthermore, no pilin-like candidates have been identified for any of the T4BSSs to date. Therefore, what these different extracellular densities are and if they are related to released/secreted proteins (such as DotA or IcmX (47)) remain unknown.

### Inner membrane stacking, a characteristic feature observed only in SCVs

In the day 15 axenic culture, 19% of cells were SCVs, 37% were LCVs and 44% are in their transition state (SI Fig. 1). The SCVs, which are 600 nm or shorter, exhibited relatively dense periplasmic space and condensed DNA compared to LCVs (SI Fig. 2). Another interesting feature in the cryotomograms of SCVs was the presence of a stack of tightly packed membranes in the cytoplasm (Fig. 3A-E, SI Movie 1). In 58 cryotomograms, we always observed a single membrane stack per SCV with each stack having 2-6 layers consistently spread ~10 nm apart as revealed by density profile analysis (Fig. 3F, SI Movie 1). This membrane stacking was generated by invaginating the IM towards the cytoplasm and was initiated in LCVs, but densely packed mature stacks were exclusively found in SCVs which suggests that maturation of these stacks is developmentally linked to the LCV to SCV differentiation.

To investigate if the membrane stacking is any different during an infection condition, we imaged the 28 days post-infection host (Vero cells)-derived *C. burnetii* variants using cryo-ET. In this sample, ~35% of the cells were SCVs, 41% were in transition state and 24% were LCVs. Similar to the day 15 axenic culture, only host-cell derived SCVs had densely packed membrane stacks while LCVs and transition state cells only revealed early stages of these membrane invaginations. Intriguingly, only in the host-cell-derived SCVs we also observed a dark linear density with an average length of few hundred nanometers (~100-300 nm) nm and located ~10 nm from the IM (SI Fig. 7). This linear density appeared morphologically rather different from the IM stacks suggesting that it is of a different nature.

### Cytoplasmic spore-like structures in SCVs and transitional cells

Another prominent feature in *C. burnetii* SCVs of the day 15 axenic culture is the presence of IM invaginations that extended and curved to form a multilayered spherical structure in the cytoplasm. Initially, the IM invaginated into the cytoplasm forming a septum with a protein-like layer visible at one side (Fig. 4 and SI Movie 2). This protein-like layer was located ~7-8 nm from the invaginated membrane. Subsequently, this septum elongated and curled with the protein-like layer still visible at one side to ultimately form a concentric multilamellar spherical cytoplasmic structure with an average diameter of ~ 120-150 nm (Fig, 4, SI Fig. 8, Movie 2). In total, we identified 16 such structures in 58 cryo-tomograms.

It was difficult to determine how many layers this structure has at the resolution of our cryo-tomograms but the number of membrane layers in these structures was variable in different cells as they had different thickness. It was also not possible to determine whether they were connected to the IM or not. In 5 examples, this structure adapted an ellipsoidal shape instead of a spherical one. This structure had polar cellular localization in 12 examples, while the remaining 4 were located at the center of the cell and appeared darker than the polar one (SI Fig. 9). In one cell, two such structures were present while the other examples had one per cell. Finally, the SCVs of the day 15 axenic culture were also associated with many outer membrane vesicles (OMVs, SI Fig. 9). Both the cytoplasmic multilayered structures and the OMVs were lacking in SCVs of the other samples, namely, the 5 day axenic culture and the host-cell-derived SCVs.

### Cytoplasmic tubes in SCVs

Another important morphological feature in the cells of the 5 day axenic culture, both at pH 7.2 and pH 4.75, was the presence of cytoplasmic tubes that were ~ 10 nm wide and up to few hundreds of nanometers long (SI Fig. 10). We identified these tubes in 43 cells in total and each cell contained many tubes, which were either single or stacked together as a bundle (SI Fig. 10). We could only identify two cells with such tubes in the host-cell-derived SCVs, and none in the day 15 axenic culture SCVs. Interestingly, these tubes were located at the center of the cell, amidst the DNA density, and were not found in areas outside the DNA where the ribosomes are present.

## DISCUSSION

Our cryo-ET imaging of the unique biphasic developmental cycle of *C. burnetii* enabled us to visualize the morphological changes associated with transition from SCV to LCV and *vice versa* at molecular resolution under native (hydrated, unfixed) conditions. In addition, it also allowed us to visualize the structural details of the T4BSS molecular machine and its developmental regulation with respect to the biphasic cell cycle.

Earlier studies on the *L. pneumophila* T4BSS revealed that these molecular machines are primarily localized at the bacterial cells poles and their polar localization is essential for *L. pneumophila* virulence (42,43). Similarly, *C. burnetii* T4BSS are also primarily located at the cell poles, but we noticed some particles away from the cell pole suggesting the polar localization in *C. burnetii* is not as tightly controlled. The two key proteins, DotU and IcmF, that target the *L. pneumophila* T4BSS to the cell poles (43), have homologues in *C. burnetii*; however, *icmF* has a point mutation which leads to a premature stop (44), which could explain why some of the T4BSS particles were away from the cell pole in *C. burnetii*.

Like *L. pneumophila*, some T4BSS particles in *C. burnetii* were also seen near the division plane suggesting that these nanomachines start assembling near the future cell pole at the beginning of the septation process. Interestingly, the T4BSS complexes were only observed in replicative LCVs and never in SCVs, which is consistent with earlier studies suggesting that the expression and activity of the *C. burnetii* T4BSS are tightly regulated with respect to developmental transitions (20–24). Moreover, the absence of T4BSS in SCVs implies that they are not essential for host-entry and early stages of infection.

Some *C. burnetii* T4BSS particles were associated with extracellular proteinaceous densities that were located right at the gate of the T4BSS particles and it remains unclear what these densities are. Earlier studies reported that the *C. burnetii* T4BSS release/secrete two of its components, namely, DotA and IcmX in a T4BSS-dependent manner when grown in host free axenic media (47) and one possibility for these extracellular densities is that they are related to such release/secretion. Intriguingly, we also noticed T4BSSs associated with extracellular short tubular structures; however, some of these tube-like structures lacked T4BSS particles beneath them. While T4ASS is known to harbor extracellular pili containing VirB2 and VirB5 proteins (48), the T4BSS has no putative pilin candidates and there are no reports of this system producing a pilus. Previous studies proposed that the *C. burnetii* T4BSSs interact with the CCV membrane using short tethers that could facilitate translocation of T4BSS effectors (49), and a contemporaneous study also reported short filamentous densities between the bacterial OM and the CCV membrane (46). It is possible that the short tubular structures above some T4BSS particles are used to establish such an intimate contact between the bacterial OM and CCV membrane.

Our *in situ* structural analysis of the *C. burnetii* T4BSS using subtomogram averaging revealed that it is architecturally similar to that of the *L. pneumophilla* T4BSS at our resolution (43,50). Recently, Sheedlo *et al*. used cryo-EM single particle analysis to resolve a near-atomic resolution structure of the *L. pneumophilla* T4BSS core complex (45). Fitting their structure into our subtomogram average allowed us to tentatively assign certain components of the OM core complex and the periplasmic ring complex. Additionally, while Sheedlo et al identified multiple species-specific proteins for the *L. pneumophilla* T4BSS - Dis1 (Lpg0657), Dis2 (Lpg0823), and Dis3 (Lpg2847) (45), the counterparts of those proteins, if any, in the *C. burnetii* T4BSS remain unknown.

Another important morphological feature associated with the biphasic developmental cycle of *C. burnetii* is the presence of tightly packed membrane layers directly below the IM in the day 15 axenic culture and in host cell derived (28 dpi) SCVs. Similar membrane invaginations were reported by McCaul *et al* nearly 4 decades ago using traditional TEM of chemically fixed SCV cells (4). These membrane stacks could be a way to store membranes for future rapid developmental transitions to LCVs. Interestingly, the day 15 axenic culture cells also produced many OMVs, not seen in other samples, which could be a result of depleted nutrients in the medium.

A controversial point in the *C. burnetii* biology is related to whether it can produce spores. Conventional TEM experiments of fixed and negatively stained *C. burnetii* cells, performed since 1980s, indicated the presence of small multilayered structures in the periplasm of some cells (51). Based on morphological similarity between these structures and the spores of other bacterial species, it was hypothesized that LCVs could produce endogenous spores and ultimately release them into the environment with other SCV cells where the spores eventually also transform into SCV to invade new host cells (51). Additionally, it was shown that these structures contain DNA, and a homologue of the sporulation gene SpoIIIE was identified in *C. burnetii* (52,53). Subsequent studies reported these supposed “endospores” also in *C. burnetii* cells purified from the heart valves of patients diagnosed with Q fever endocarditis (54).

However, the nature of these structures remained controversial for a long time [for a nice review see (55)], hence they are usually referred to as spore-like particles (SPL) [See references (55,56) and references therein]. There are multiple reasons now to state that sporulation does not happen in *C. burnetii* including the absence of the biochemical spore marker, dipicolinic acid (57), and more importantly, the sequencing of *C. burnetii* genome revealed that it lacks sporulation genes (58). Our results here are also in accordance with this notion. First, these structures have different shapes, sizes, layers, and cellular localization which is inconsistent with a tightly regulated sporulation process. Second, we never detected these structures independently outside cells and only observed them inside cells (grown in ACCM2 medium for 15 days).

Finally, SCVs from the axenic culture day 5 (both at pH 4.75 and pH 7.2) contained bundles of cytoplasmic tubes ~10 nm in diameter amidst the DNA region of the cell. Morphologically, they are reminiscent of filamentous bundles of unknown functions described in other bacterial species (59). The fact that these bundles are always present inside the DNA region in the cell suggests they could be related to other protein-DNA polymers such as those formed by RecA and MuB, which are also ~ 9.5 nm *in vitro* (60,61). Another possibility is that these are a protective crystalline form of DNA and DNA-binding proteins known to form under stress conditions (62). The other two interesting uncharacterized morphological features include the linear density seen under the IM in host-derived SCVs *C. burnetii*, and the protein-like polymer seen on the side of early IM invaginations, which might play a role in the formation of the membrane stacks and/or spherical multilayered structures. Further studies are required to understand the biogenesis and actual function of the multilayered membranes stacks, ‘endo-spore’ like structures and other uncharacterized features described in this study.

## MATERIALS AND METHODS

### Strains and growth conditions

The *Coxiella burnetii* Nine Mile phase II (clone 4, RSA 439) strain was used. For samples purified from eukaryotic host cells, *C. burnetii* was grown in African green monkey kidney (Vero) fibroblasts (CCL-81; American Type Culture Collection) which are cultured in RPMI medium (Invitrogen, Carlsbad, Calif.) supplemented with 2% fetal bovine serum as described in reference (18). Briefly, Vero host cells were infected in 150-cm^2^ flasks for 4 weeks without changing the growth medium. During the first week, cells were incubated at 37°C in 5% CO_2_ and subsequently the flasks containing the cells were kept for three weeks at room temperature with their caps tightened. Bacterial cells were then purified from the infected eukaryotic cells using Renografin density gradient centrifugation (63). Purified SCV cells were resuspended in K-36 buffer (0.1 M KCl, 0.015 M NaCl, 0.05 M potassium phosphate, pH 7.0) and stored at −80°C until they were thawed before plunge-freezing as described below.

Alternatively, *C. burnetii* (wild type Nine Mile phase II strain or an isogenic *dot/icm* mutant of this strain) was grown under axenic cultivation conditions as described in (64,65). To cultivate *C. burnetii*, ACCM-2, which is a modified ACCM medium, was used (64). In this modified medium, 1 mg/ml methyl-β-cyclodextrin (Mβ-CD) is used to replace 1% FBS. Bacterial cells were cultured in this medium in T-75 flasks or 0.2-μm-pore-size-filter-capped 125-ml Erlenmeyer flasks with 20 ml of medium, T-25 flasks with 5 ml of medium, or 3 ml of medium in each well of a 6-well tissue culture plate. Bacterial cultures were kept in a CO-170 incubator (New Brunswick Scientific, NJ) with the following conditions: 37° C in a 2.5% O_2_ and 5% CO_2_. Subsequently, cells were washed and resuspended either in PBS pH 7.2 or in Citrate buffered saline pH 4.75 (PBS + 10 mM citrate) and then frozen at −80°C until they were thawed before plunge-freezing as described below.

### Cryo-ET sample preparation and imaging

R2/2 carbon-coated 200 mesh copper Quantifoil grids (Quantifoil Micro Tools) were first glow-discharged for 60 seconds. 1 μL of BSA-treated 10-nm gold solution was added to 4 μL of cells thawed from a −80°C stock. The combination of cells and gold was then pipetted on the grids in a Vitrobot chamber (FEI) with 100% humidity. The extra fluid was blotted off using a Whatman filter paper and the grids were plunge-frozen in a liquid ethane/propane mixture. The samples were imaged using an FEI Polara 300 keV field emission gun electron microscope (FEI, Hillsboro, OR, USA) equipped with a Gatan image filter and K2 Summit direct electron detector in counting mode (Gatan, Pleasanton, CA, USA). Data were collected using the UCSF Tomography software (66) with each tilt series ranging from −60° to 60° in 1°, 2° or 3° increments, an underfocus of ~7 μm, an electron dose of 130 e^-^/Å^2^, and a pixel size of 3.9 Å.

### Image processing and subtomogram averaging

Three-dimensional reconstructions of tilt-series were done with either IMOD software package (67) or using EMAN2 software (Chen et al., 2019). In total, 88 tomograms of *C. burnetii* grown at pH 7 and 69 tomograms of *C. burnetii* grown at pH 4 were of sufficient quality for further sub-volume analysis. Particles were manually identified and selected from tomograms in each dataset. In the tomograms of *C. burnetii* grown at pH 7, 726 particles were identified, and 414 particles were identified in tomograms of *C. burnetii* grown at pH 4. Particles were extracted using a box size of 96 from tomograms at x2 binning (8.06 Å/pix). The EMAN2 program e2spt_sgd_new.py was used to generate initial models using a subset of 60 high-quality particles from each dataset at 4x binning. Initial models with C1 symmetry were generated with default parameters in the e2spt_sgd_new.py program. For initial models with C13 symmetry, the T4SS C1 initial models were aligned to the symmetry axis and used as references. Initial models with C1 or C13 were filtered to 50 Å resolution and used for subsequent sub-tomogram refinement. Extracted particles from each dataset were used for sub-tomogram averaging with either C1 or C13 symmetry. Particles with x2 binning were subjected to two rounds of 3D particle orientation refinement, 2D sub-tilt translation refinement, and sub-tilt translation & rotation refinement, followed by a sub-tilt defocus refinement. Finally, soft masks for either the Outer membrane core complex (OMCC) or Inner membrane were applied to focus the alignment of particles to each respective region.

The density projection profile of the cell envelope and membrane invaginations was calculated in ImageJ (Schindelin et al., 2012) using a 50 nm wide linear section of the cell envelope. Density plots were generated using Microsoft excel software.

To improve segmentation quality selected tomograms were missing wedge corrected using IsoNet (68). Membranes, membrane invaginations, and spores were manually segmented using drawing tools in IMOD (69). Outer membrane vesicles were segmented with a convolutional neural network in EMAN2 (70). Alternatively, deep convolutional neural networks (CNNs) were trained using the manually generated ground truth and fine-tuned until a good quality of segmentation was reached for automated segmentation in 3D. The segmentation has been proofread and cleaned-up by expert inspection using the open-source software Knossos (https://knossos.app/). For each segmentation class, meshes and binary Tiff masks were generated for subsequent rendering and visualization in Knossos and in the open-source 3d visualization tool Blender (https://www.blender.org/). Data visualization was also performed using ChimeraX (71).

## Acknowledgements

This project was funded by the National Institutes of Health (grant R01 AI127401 to G.J.J), an NHMRC grant (APP1196924 to DG), and the Intramural Research Program of the National Institutes of Health, National Institute of Allergy and Infectious Diseases (Grant number AI000931-20 to R.A.H. and C.L.L.). DCS is supported by the Melbourne Research Scholarship. We are grateful to Prof. Hayley Newton for critically reading the manuscript, Prof. Elitza Tocheva for insightful discussions, and Somavally Dalvi for her help with figure preparation.

## Supporting Information

**SI Fig. 1.**
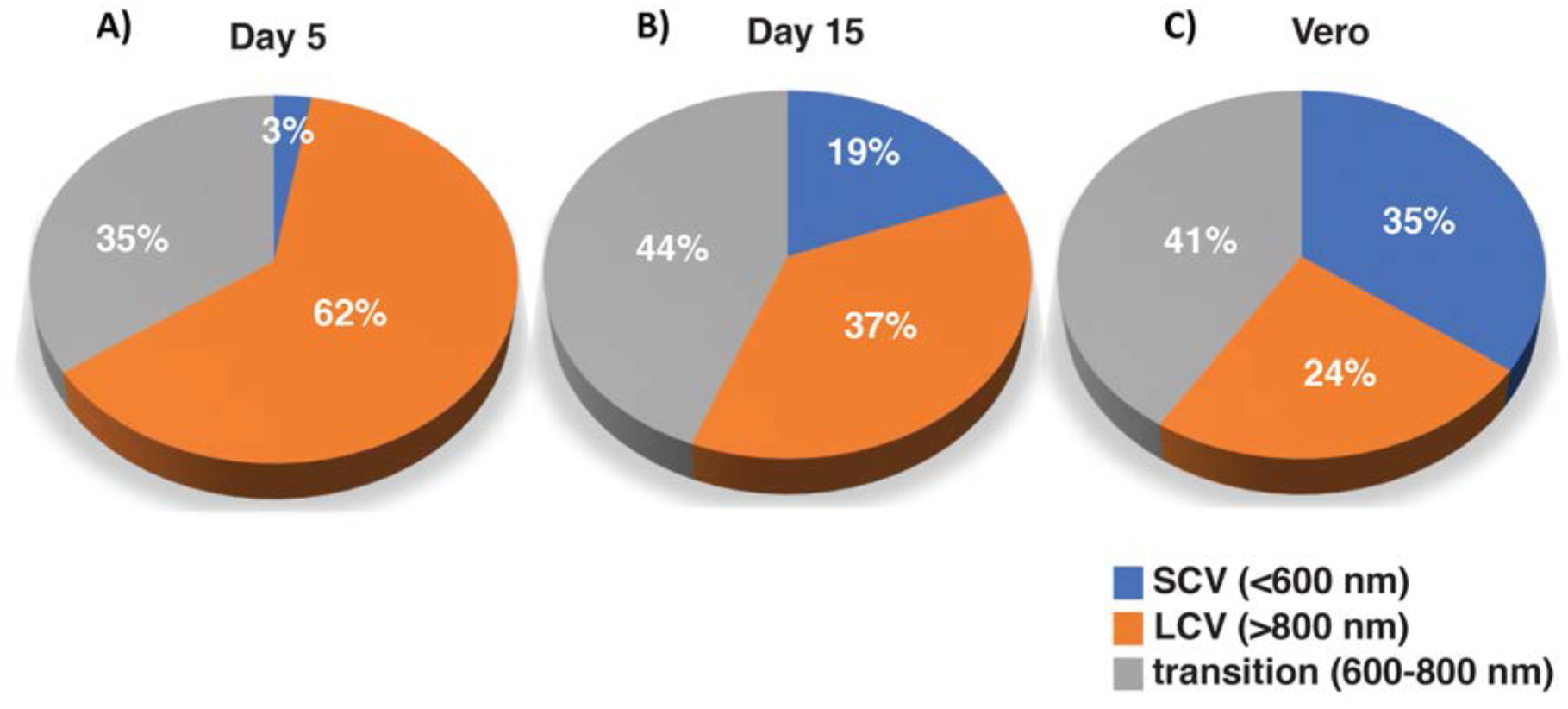
Distribution of *C. burnetii* SCVs, LCVs and transition state variants (TSVs) in axenic culture and under infection conditions. *C. burnetii* variants in axenic culture (A) day 5, (B) day 15. C) *C. burnetii* cells purified from infected Vero cells, 28 dpi. SCVs (<600nm), LCVs (>800nm), and TSVs (600-800nm). At least 70 individual *C. burnetii* cells lengths were inspected for different conditions.

**SI Fig. 2.**
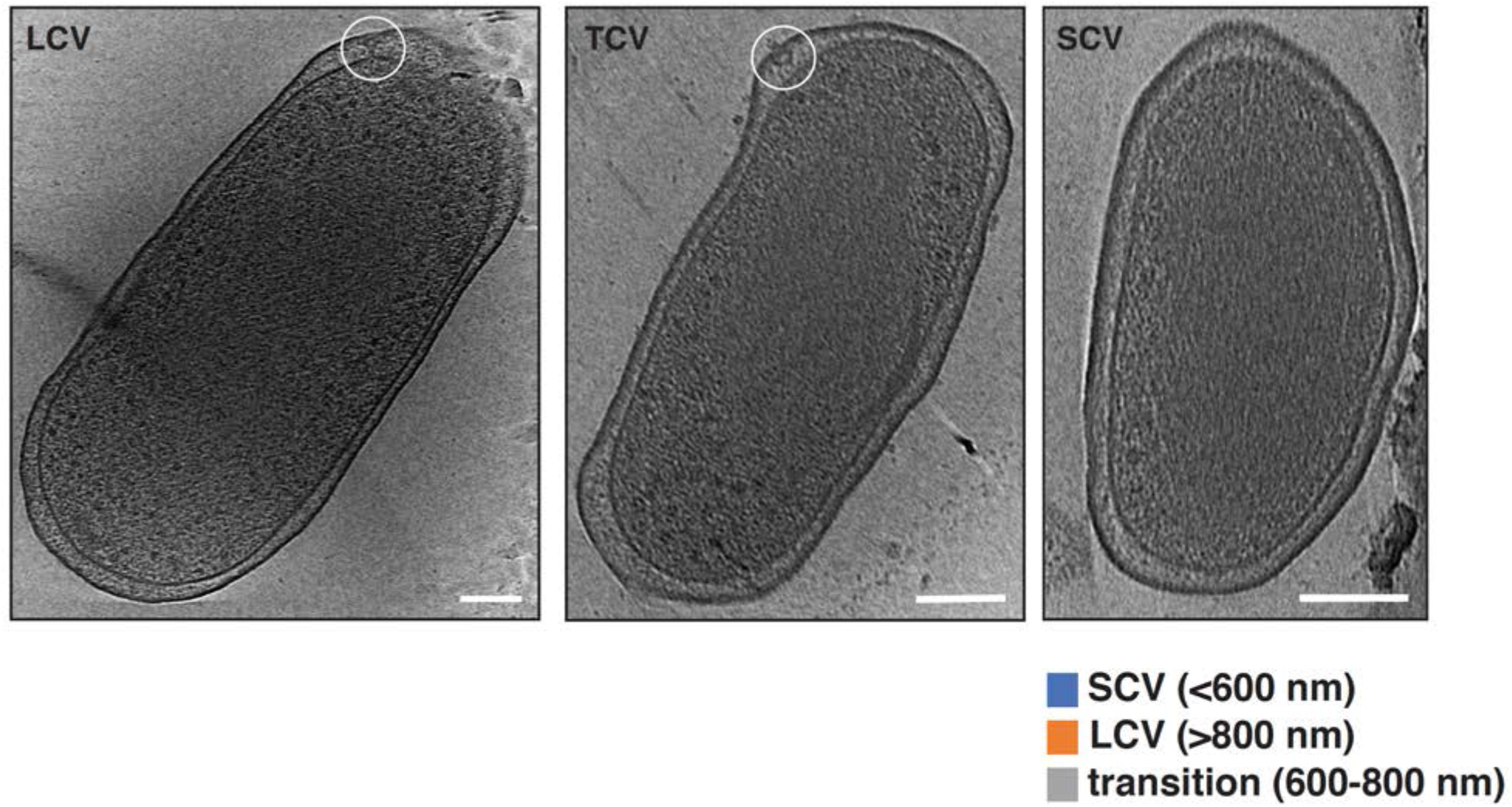
Tomographic slices of *C. burnetii* cells (LCV, TSV and SCV) highlighting the presence of T4BSS particles (white circles) in the polar region of LCV and TSV or absence the T4BSS in SCV. Scale bar 100 nm.

**SI Fig. 3.**
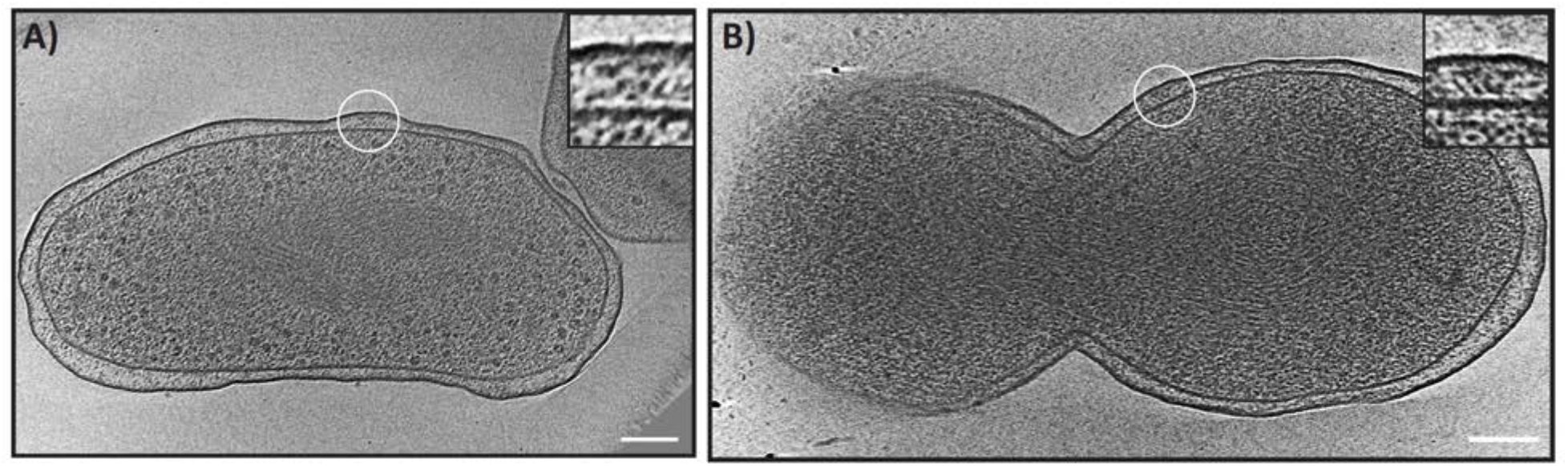
(A-B) Tomographic slices of *C. burnetii* cells showing the presence of T4BSS particles (white circles) away from the cell pole (A) and near the division site (B). Scale bar 100 nm. Insets showing enlarged views of the T4BSS particles.

**SI Fig. 4.**
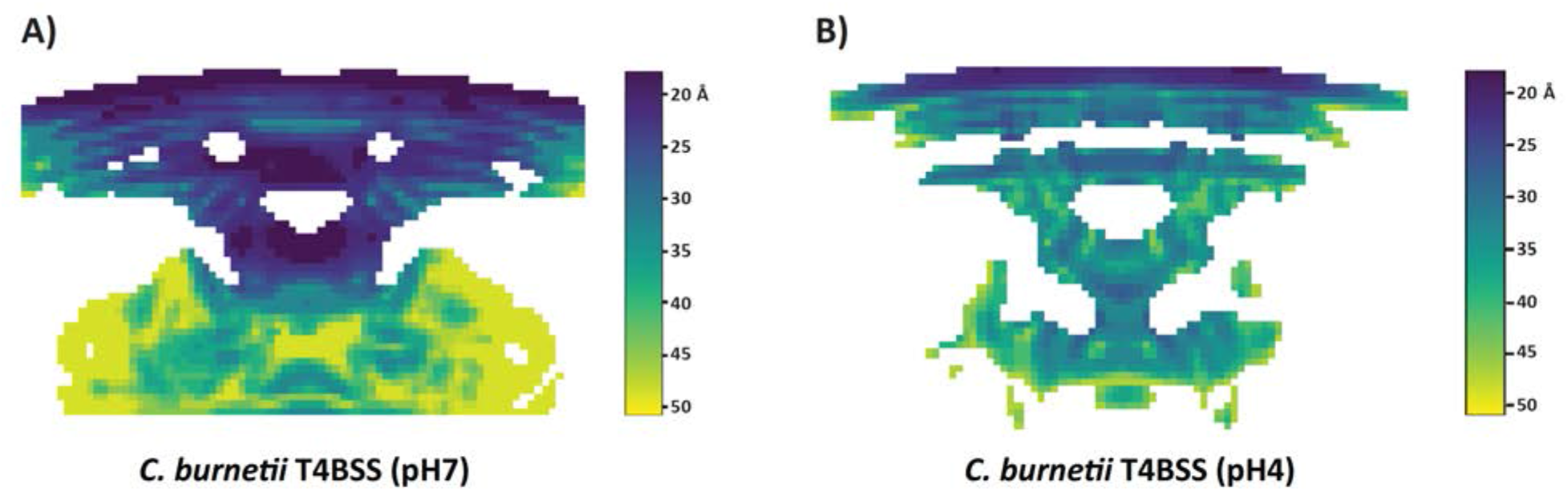
Local resolution of the *C. burnetii* T4BSS subtomogram averages calculated by ResMap. A) *C. burnetii* T4BSS average at pH 7 and (B) *C. burnetii* T4BSS average at pH 4.

**SI Fig. 5.**
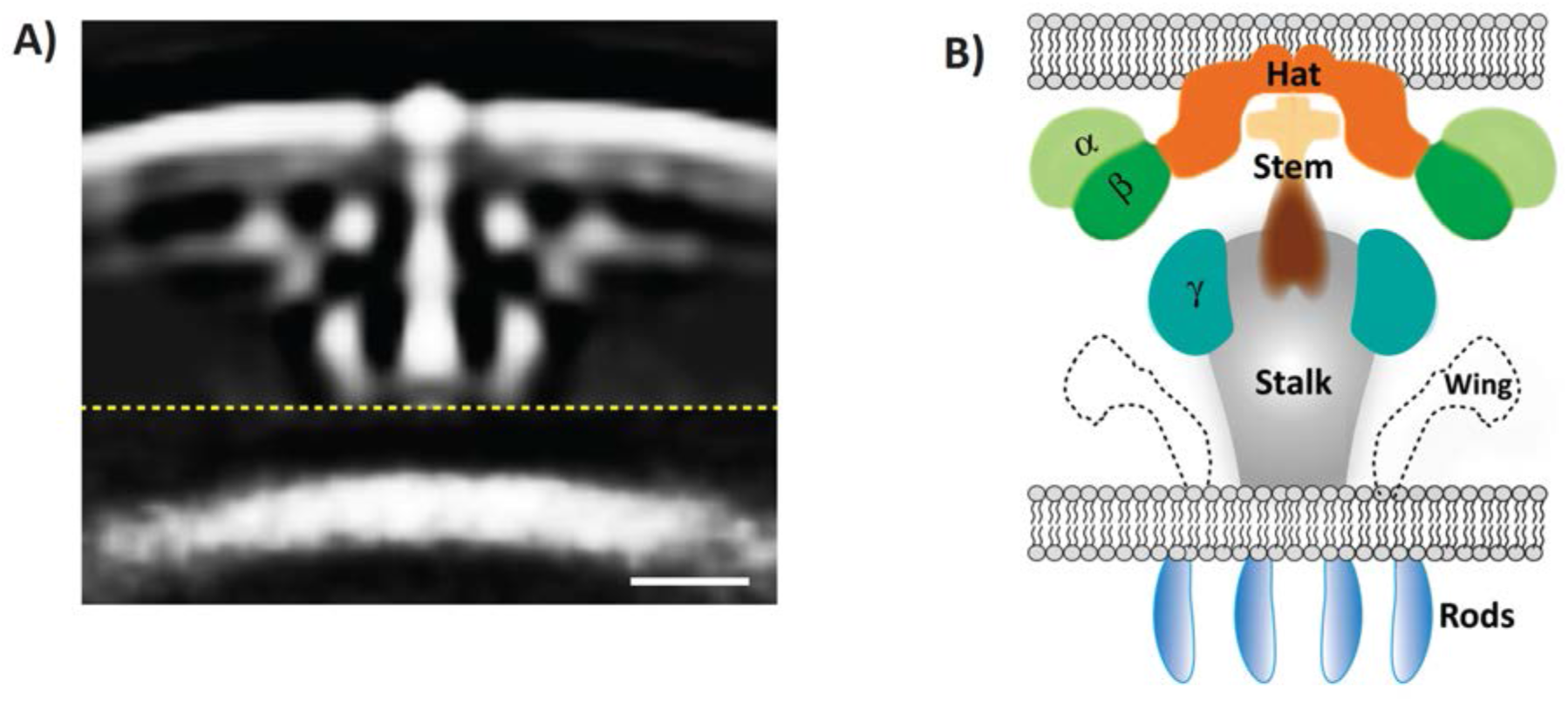
(A) A central slice through the subtomogram average of the T4BSS of *C. burnetii* at pH7. Dashed yellow line indicates a composite of two averages obtained with different masks (see materials and methods) concatenated together. Scale bar 10 nm. (B) A schematic of the *L. pneumophila* T4BSS showing distinct densities (alpha, beta, hat, stem, gamma, stalk, wing and ATPases/rods) as reported by Ghosal et al (37).

**SI Fig. 6.**
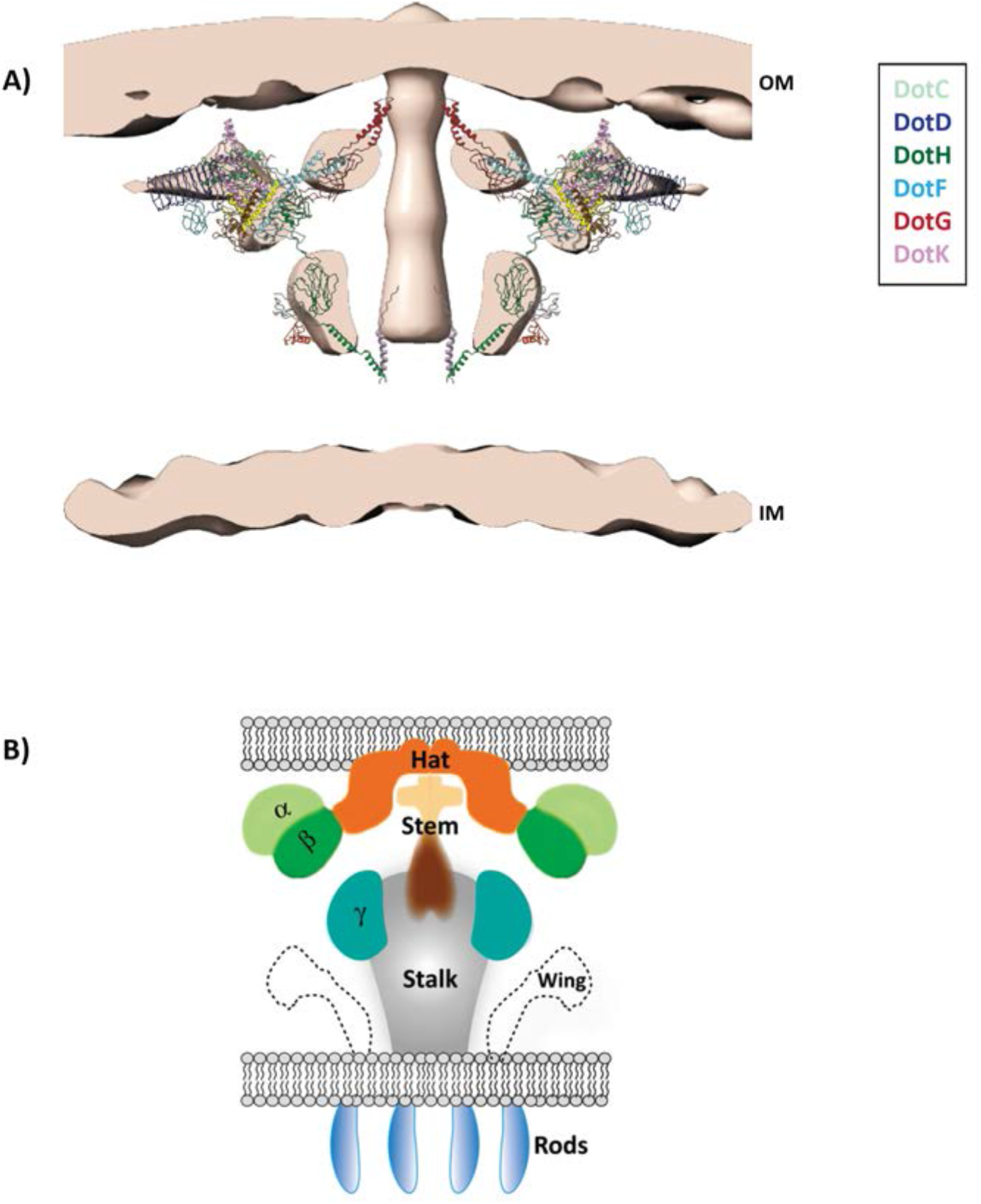
An atomic models of the *L. pneumophila* T4BSS core complex (45) fitted into our *C. burnetii* T4BSS subtomogram average (day 5 axenic culture, reproduced from Fig. 1). In the atomic model, DotC (light green), DotD (midnight blue), DotH (green), DotF (corn flower blue), DotG (maroon), DotK (pink) constitute the outer membrane complex. B) A schematic of the *L. pneumophila* T4BSS showing distinct densities (alpha, beta, hat, stem, gamma, stalk, wing and ATPases/rods) as reported by Ghosal et al (37).

**SI Fig. 7.**
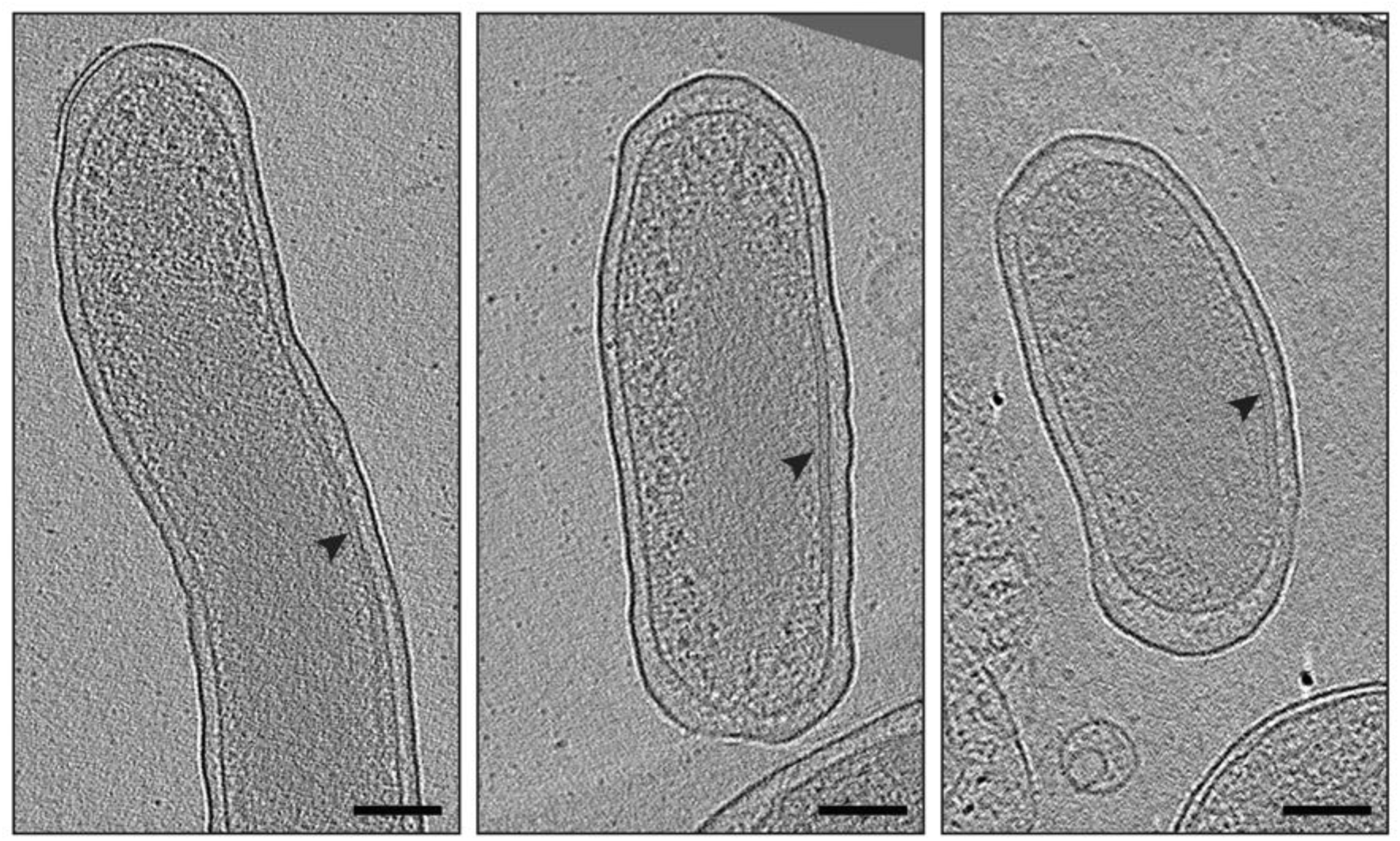
Tomographic slices of *C. burnetii* cells derived from infected Vero cells (28 dpi) highlighting the presence of a thin line (arrowheads) just below the IM. Scale bar 100 nm.

**SI Fig. 8.**
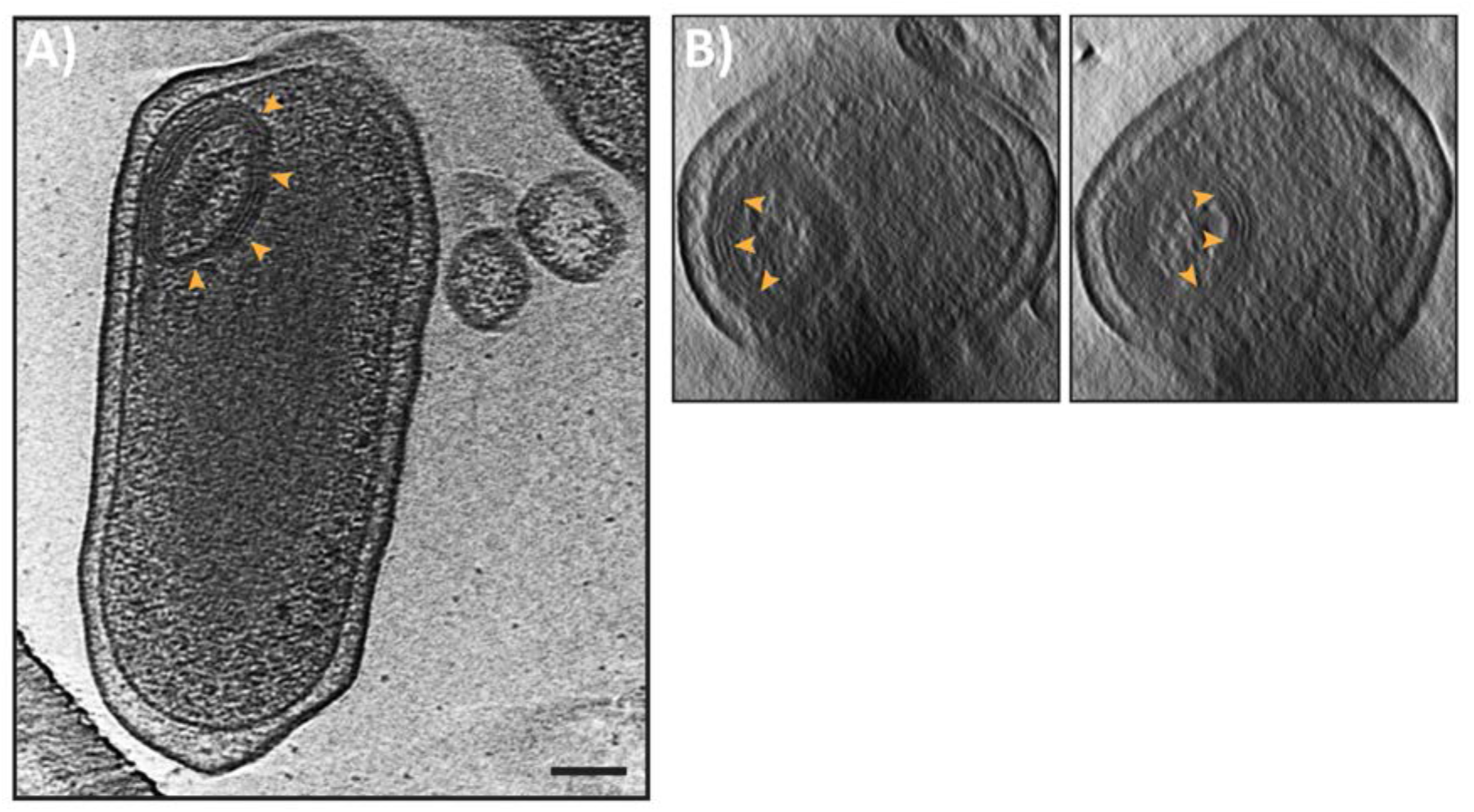
A) Tomographic slice through a *C. burnetii* cell (axenic culture, day 15) showing concentric multilamellar structure just below the IM (orange arrowheads). B) End on views of the same cell showing these structures are spherical (orange arrowheads). Scale bar 100 nm.

**SI Fig. 9.**
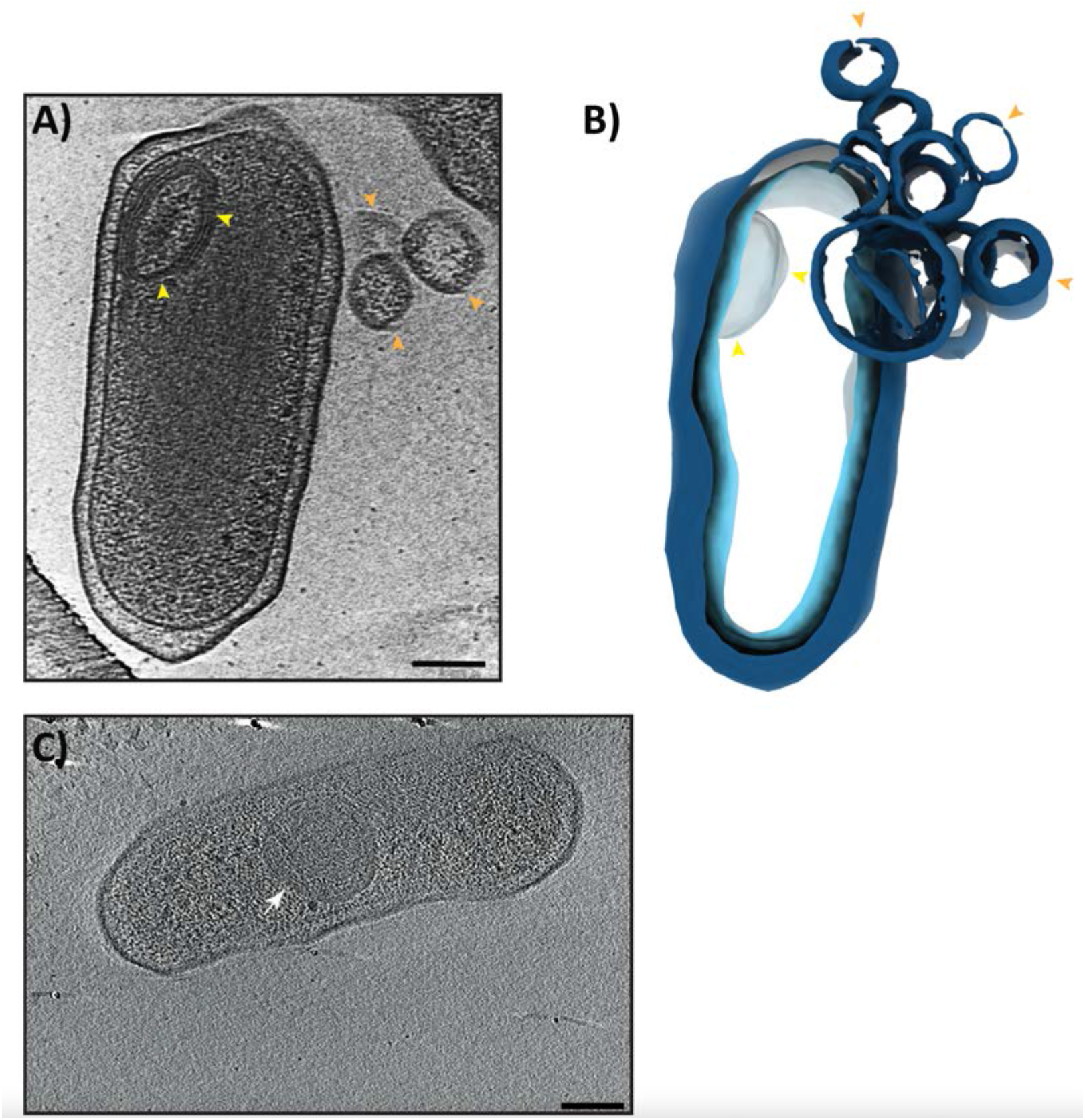
A) Slice through an electron cryotomogram of a *C. burnetii* cell (axenic culture, day 15) showing concentric multilamellar spherical cytoplasmic polar structure (yellow arrowheads) and outer membrane vesicles (orange arrowheads). B) A segmented volume of the same cell shown in (A) highlighting the multilamellar spherical cytoplasmic structure (yellow arrowheads) and OMVs (orange arrowheads). C) A Slice through an electron cryotomogram of a *C. burnetii* cell indicating the presence of concentric multilamellar spherical cytoplasmic central structure (white arrow). Scale bar 100 nm.

**SI Fig. 10.**
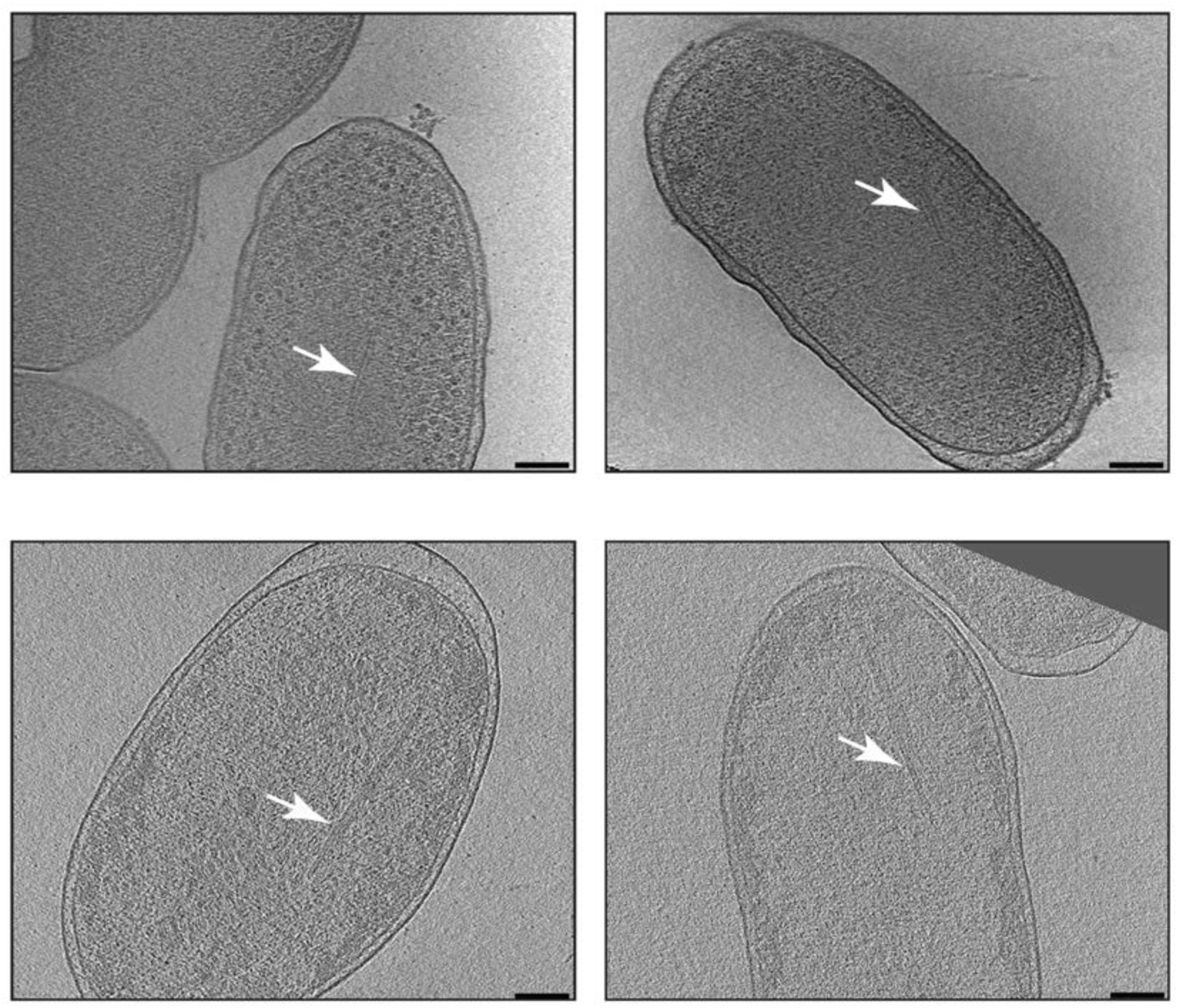
Tomographic slices of *C. burnetii* cells grown in axenic culture day 5 pH 7.2 (top panels) and pH 4.75 (bottom panels) highlighting the presence of a thin tubular filaments (white arrows) in the cytoplasm. Scale bar 100 nm.

### Supplementary Information Movie legends

**SI Movie 1:** Three-dimensional (segmented) view of a host-derived (28 dpi) *C. burnetii* cell showing a stack of a membrane associated with the IM.

**SI Movie 2:** Three-dimensional (segmented) view of a *C. burnetii* cell grown in axenic culture (day 15) showing concentric multilamellar spherical cytoplasmic structure just below the IM.

